# ST6GAL1 sialyltransferase promotes acinar cell survival and tissue regeneration during pancreatitis

**DOI:** 10.64898/2026.09.26.754701

**Authors:** Michael P. Marciel, Nikita Bhalerao, Barnita Haldar, Sejal S. Shinde, Joshua C. Anderson, Christopher D. Willey, Srikanth Iyer, Vikas Dudeja, Susan L. Bellis

## Abstract

The role of aberrant glycosylation in pancreatitis remains an under-investigated area of research. Here, we determined that the ST6GAL1 sialyltransferase, which adds α2-6-linked sialic acids to *N*-glycoproteins, was upregulated in pancreatic tissues from patients with acute (AP) and chronic (CP) pancreatitis, and in mice with experimental AP. Within these tissues, ST6GAL1 was selectively expressed in acinar cells undergoing acinar to ductal metaplasia (ADM), a process whereby injured acinar cells de-differentiate and re-enter the cell cycle to enable tissue repair. To study the functional role of ST6GAL1 in pancreatitis, we used HPNE pancreatic epithelial cells with modulated ST6GAL1 expression, along with pancreatic organoids from mice with transgenic expression of ST6GAL1 (“SC” mice). In these models, ST6GAL1 activity promoted the activation of EGFR (a well-known ADM-driver), ERK, and AKT. ST6GAL1-expressing cells also displayed an ERK-dependent upregulation of the anti-apoptotic proteins, Mcl-1 and Bcl-xL, and impaired apoptotic signaling by the TNFR1 and Fas death receptors. Unbiased kinomics profiling revealed that ST6GAL1 induced the activation of many receptor tyrosine kinases associated with cell survival and proliferation (e.g., EGFR, PDGFR, MET). Furthermore, ST6GAL1 enhanced the survival and proliferation of cells exposed to stress-inducing conditions such as serum withdrawal. Based on these findings, we hypothesized that the pro-survival phenotype imparted by ST6GAL1 would facilitate tissue healing following a bout of pancreatitis. Accordingly, we induced AP in wild type and SC mice via L-arginine injection and found that SC mice had more rapid and efficient tissue repair, evidenced by accelerated restoration of the acini, diminished acinar apoptosis, and reduced immune cell infiltration. Together, these findings highlight a novel glycosylation-dependent mechanism that drives cell survival, positioning ST6GAL1 as a key adaptive mediator of pancreatic regeneration.

## Introduction

Pancreatitis is an inflammatory disorder whereby pancreatic parenchyma is damaged and replaced by fibrous tissue, leading to pain, diabetes, and malnutrition (1, 2). Pancreatic acinar cells are injured as part of this process, often culminating in acinar cell death. However, a subset of acinar cells responds to tissue injury by de-differentiating into ductal-like, progenitor cells in an event known as acinar-to-ductal metaplasia (ADM) (3, 4). Cells undergoing ADM re-enter the cell cycle, allowing for proliferation and replacement of damaged acini (3, 4). Accordingly, ADM-like cells serve as “facultative progenitors” which are crucial for repair of the exocrine pancreas (5). Following tissue healing, ADM-like cells typically revert to their mature, quiescent acinar state. However, repeated bouts of acute pancreatitis (AP) can cause persistent inflammation, fibrosis, and continual ADM, leading to chronic pancreatitis (CP) (6). As another concern, CP is a major risk factor for pancreatic ductal adenocarcinoma (PDAC) (7), the most common form of pancreatic cancer. Cells in an ADM state are more susceptible than mature acinar cells to neoplastic transformation by the KRas oncogene (4), which is mutated in >90% of PDAC patients (8). Hence, ADM is considered the first step toward pancreatic oncogenesis.

One of the principal molecular drivers of ADM is the EGFR receptor tyrosine kinase (RTK) (9–12). EGFR is activated in acinar cells by its ligands EGF and TGFα, which are greatly upregulated within the pancreatitis microenvironment (13, 14). The importance of EGFR in acinar plasticity is underscored by the fact that acinar cells in mice with transgenic expression of TGFα spontaneously undergo ADM (9, 10). In the canonical EGFR pathway, ligand binding induces EGFR dimerization and autophosphorylation, triggering several downstream signaling cascades such as MEK-ERK and PI3K-AKT. These pathways are key mediators of cell survival and proliferation. Not surprisingly, EGFR signaling is a central player in the regeneration and repair of injured tissues in multiple organs including the pancreas (15–19).

While the mechanisms governing EGFR activation are of intense research interest, the role of variant glycosylation on EGFR structure and function remains poorly understood. We and others have shown that EGFR is a substrate for the ST6GAL1 sialyltransferase (20, 21), a Golgi enzyme that adds α2-6-linked sialic acids to select *N*-glycoproteins destined for the plasma membrane or secretion (22, 23). The α2-6 sialylation of EGFR enhances its activity, leading, in turn, to the activation of ERK, AKT, and other signaling mediators (24, 25). In cancer cells, the sialylation of EGFR promotes migration and invasion (21, 25), epithelial to mesenchymal transition (EMT) (21), and resistance to therapeutic EGFR inhibitors including cetuximab (20). Mechanistically, the α2-6 sialylation of EGFR facilitates receptor dimerization, autophosphorylation, and higher order clustering, resulting in increased downstream signaling (24). EGFR sialylation also promotes receptor recycling to the cell surface while inhibiting lysosomal-mediated degradation (24). In tandem with its effects on EGFR, ST6GAL1 sialylates other critical receptors that govern cell fate such as the Fas and TNFR1 death receptors. The ST6GAL1-mediated sialylation of Fas and TNFR1 prevents apoptosis by hindering ligand-induced Fas and TNFR1 internalization (26–28), an event required for caspase activation (29). Remarkably, these effects are specific to the α2-6 sialic acid linkage, as α2-3 sialylation does not suppress TNFR1/Fas-mediated apoptosis (28). The ligands for Fas and TNFR1, FasL and TNFα, respectively, are elevated within an inflammatory milieu, and indeed, TNFα directly contributes to the pathogenic features of pancreatitis (30). The combined actions of ST6GAL1 on receptors such as EGFR, Fas, and TNFR1 position ST6GAL1 as a pivotal cell survival factor.

ST6GAL1 plays well-established roles in malignancy. ST6GAL1 expression is dramatically increased in PDAC and other cancers, in part, through transcriptional activation induced by KRas and other oncogenes (23, 31). High ST6GAL1 expression correlates with elevated tumor stage and grade (32–35), tumor invasiveness (36, 37) and poor clinical outcomes (37–41). Conversely, ST6GAL1 expression is very low in many normal epithelial cell populations including pancreatic acinar cells (32, 38). To interrogate oncogenic functions for ST6GAL1 in PDAC, our group previously generated a mouse model with transgenic expression of ST6GAL1 in the pancreas, referred to as “SC” mice (32). SC mice were crossed to the “KC” PDAC mouse model (42), which expresses an activated KRas mutant, KRas^G12D^, in the pancreas. Mice with dual expression of ST6GAL1 and KRas^G12D^, termed “KSC” mice, had greatly accelerated PDAC development, metastatic progression and mortality when compared with KC mice (32). In this same study, ST6GAL1 was found to fuel neoplasia through facilitating acinar transition into an ADM-like state. Acinar cells from SC mice (i.e., with wild-type KRas) displayed a de-differentiated phenotype characterized by the downregulation of mature acinar genes (*Ptf1a)*, upregulation of ductal genes (*Sox9*, *Krt8*, *Krt9*), and a pronounced activation in stemness signaling networks including the Wnt and Notch pathways. These findings align with other reports suggesting an association between ST6GAL1 and stemness. ST6GAL1 is a marker for several stem/progenitor populations including neural progenitor cells (43), gastric stem cells (44), induced pluripotent stem cells (45, 46) and cancer stem cells (38). Importantly, ST6GAL1 plays a functional role in conferring stemness properties (32, 38, 46).

Despite evidence implicating ST6GAL1 as a mediator of a progenitor-like state, the potential involvement of ST6GAL1 in tissue regeneration remains largely unexplored. In the current study, we investigated whether ST6GAL1’s pro-survival activity aids in tissue recovery following pancreatitis. We first determined that ST6GAL1 is markedly upregulated in the ADM-like acinar cells of AP and CP patient tissues, and in murine experimental AP induced by either L-arginine (L-arg) (47) or cerulein (48). This finding offers new insight into the dynamic regulation of ST6GAL1 expression within an inflammatory setting. We then evaluated the functional effects of ST6GAL1 on pancreatic epithelia using two models: (1) hTERT-immortalized <u>H</u>uman <u>P</u>ancreatic <u>N</u>estin-<u>E</u>xpressing cells (hereafter abbreviated as “HPNE” cells) engineered with or without ST6GAL1 expression and (2) pancreatic organoids derived from SC and wild type (WT) mice. In both models, ST6GAL1 activity stimulated signaling by EGFR, ERK and AKT, and induced the expression of the anti-apoptotic proteins, Mcl-1 and Bcl-xL. On the other hand, ST6GAL1 suppressed apoptosis directed by TNFR1 and Fas.

Phenotypic assays revealed that ST6GAL1-expressing cells exhibited better survival and proliferation under conditions of cell stress. Finally, we induced AP in WT and SC mice via L-arg administration and found that SC mice had more rapid and effective pancreatic tissue recovery. Taken together, these results highlight a new function for ST6GAL1 in promoting tissue regeneration following inflammatory assaults such as pancreatitis.

## Results

### ST6GAL1 expression is upregulated in CP and AP patient tissues and in murine models of AP

ST6GAL1 expression was evaluated by immunohistochemistry (IHC) in CP patient tissues displaying regions of pancreatitis-induced acinar damage, or in healthy adjacent tissue. In healthy tissues, ST6GAL1 was not detected in acinar cells (Figure 1A, asterisk), although moderate expression was noted in the islets (Figure 1A, arrowhead), as previously reported (32). In CP tissues, ST6GAL1 was selectively upregulated in acinar cells that displayed an abnormal, ductal-like morphology (Figure 1A, arrows). Quantification of the number of ST6GAL1-expressing cells in the exocrine pancreas confirmed increased ST6GAL1 levels in CP (Figure 1B). Similarly, ST6GAL1 expression was elevated in the acinar cells of patients with AP (Figure 1C). ST6GAL1 expression was then examined in two murine pancreatitis models: L-arg-stimulated AP (47) and cerulein-stimulated AP (48). In both models, ST6GAL1 expression was found in acinar cells with a ductal-like morphology (Figure 1D and E), whereas ST6GAL1 was not detected in the acinar cells of mice injected with the saline control. These results indicate that ST6GAL1 is upregulated in human and murine pancreatitis tissues.

**Figure 1.**
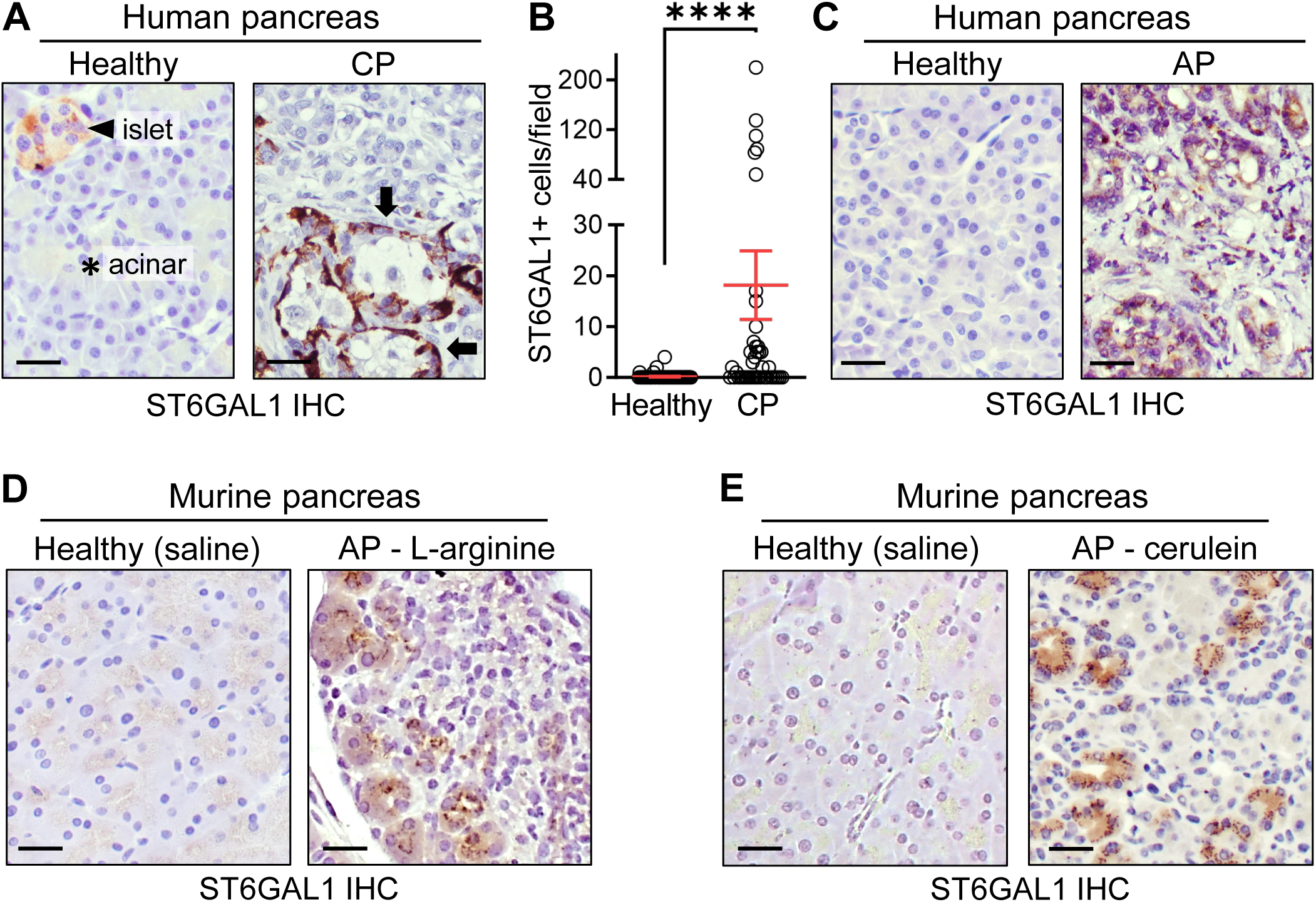
ST6GAL1 expression is upregulated in pancreatitis tissues. **(A)** ST6GAL1 IHC in pancreatitis tissues from CP patients or healthy adjacent pancreas. In the healthy pancreas, ST6GAL1 is expressed in the islets (arrowhead), but not acinar cells (asterisk). In CP tissues, ST6GAL1 is expressed in acinar cells with abnormal morphology (arrows). **(B)** Quantification of ST6GAL1-positive cells in the human exocrine pancreas. Graph depicts mean ± S.E.M., data analyzed by a D’Agostino-Pearson Test followed by a Mann Whitney test. For healthy tissue, n = 54; for CP, n = 43; **** *p* < 0.0001. **(C)** Increased ST6GAL1 expression in AP patient tissues. **(D-E)** Increased ST6GAL1 expression in AP tissues from mice injected with L-arg (**D**) or cerulein (**E**). Control groups were injected with saline. Scale bars = 50 μm; original magnification, 200×.

### Pancreatitis-induced ST6GAL1 expression is associated with a proliferative, ADM-like phenotype

The cerulein AP model was used to interrogate the effects of ST6GAL1 on acinar phenotype. Pancreata from mice injected with cerulein or saline were evaluated by immunofluorescent (IF) microscopy for the expression of: ST6GAL1; amylase (acinar marker); and Sox9 (ductal marker). The abnormal expression of Sox9 in acinar cells is a well-accepted hallmark of ADM (49, 50), and ST6GAL1 is known to promote Sox9 expression (32, 38). In the healthy acinar cells of the saline cohort, there was no detectable expression of ST6GAL1 or Sox9 (Figure 2A). However, ST6GAL1, amylase and Sox9 were co-expressed in cells of the cerulein group, confirming upregulation of ST6GAL1 in ADM-like cells (Figure 2A). Many of the ST6GAL1-positive cells in the cerulein samples co-expressed the proliferative marker, Ki67, suggesting these cells were actively cycling (Figure 2B). Tissues were also stained for the apoptotic marker, cleaved caspase 8 (cl. casp8). Many apoptotic acinar cells were identified in the cerulein cohort, however these cells did not co-express ST6GAL1 (Figure 2C). In the healthy pancreas, acinar cells typically lacked expression of Ki67 and cl. casp8.

**Figure 2.**
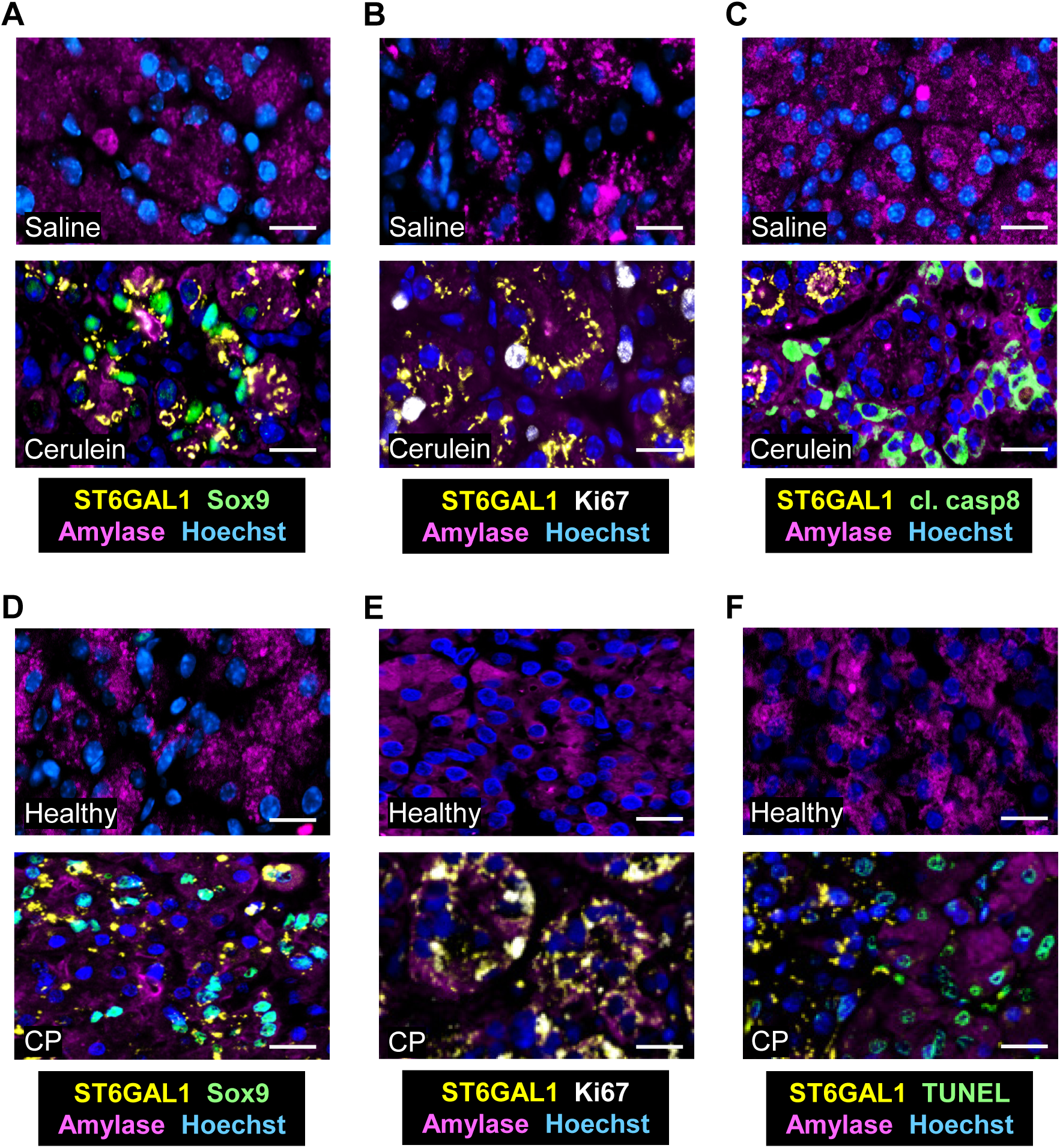
Pancreatitis-induced ST6GAL1 expression is associated with a proliferative, ADM-like phenotype. **(A-C)** Mice were injected i.p. with saline or cerulein to induce AP. **(A)** Tissues were evaluated for co-expression of Sox9 (green), ST6GAL1 (yellow), and the acinar marker, amylase (magenta). ADM-like cells (co-expressing ST6GAL1, Sox9, and amylase) were identified in the cerulein cohort but not controls. **(B)** Tissues were stained for Ki67 (white), ST6GAL1 (yellow), and amylase (magenta). Proliferative cells co-expressing ST6GAL1, Ki67, and amylase were identified in the cerulein, but not control, group. **(C)** Tissues were stained for the apoptotic marker, cleaved caspase 8 (cl. casp8, green), ST6GAL1 (yellow), and amylase (magenta). Cells undergoing apoptosis were identified in the cerulein, but not control, group. However, ST6GAL1-expressing cells and cl. casp8-expressing cells were in distinct compartments. **(D-F)** IF staining of healthy human pancreas and CP patient tissues. **(D)** Tissues were stained for Sox9 (green), ST6GAL1 (yellow), and amylase (magenta). Cells co-expressing ST6GAL1, Sox9 and amylase were identified in CP, but not healthy, tissues. **(E)** Tissues were stained for Ki67 (white), ST6GAL1 (yellow), and amylase (magenta). Proliferative, ST6GAL1-positive acinar cells were found in CP tissues, but not healthy pancreas. **(F)** Tissues were evaluated for apoptosis by TUNEL staining (green), in combination with staining for ST6GAL1 (yellow), and amylase (magenta). Cells undergoing apoptosis were identified in CP, but not healthy, tissues. However, TUNEL-positive cells in CP tissues rarely expressed ST6GAL1. Scale bars = 25 μm; original magnification, 200×.

To complement the murine pancreatitis model, tissues from CP patients were stained for ST6GAL1, amylase, Sox9, and Ki67. To measure cell death, TUNEL staining was employed. In the CP specimens, high expression of ST6GAL1 was noted in cells that co-expressed amylase and Sox9 (Figure 2D), and in cells positive for Ki67 (Figure 2E). However, cells with TUNEL staining lacked expression of ST6GAL1 (Figure 2F). In the healthy human pancreas, acinar cells did not express ST6GAL1 or Sox9, and had very low levels of Ki67 and TUNEL positivity. Quantification of the IF data from murine and human tissues revealed that ST6GAL1 was co-expressed in the majority of acinar cells with Sox9 expression, and approximately a third of the Ki67-positive cells (Table 1). Co-expression of ST6GAL1 with apoptotic markers was rare. These data suggest that the pancreatitis-induced expression of ST6GAL1 in acinar cells is associated with a proliferative and apoptosis-resistant phenotype.

**Table 1.** Quantification of IF microscopy data from pancreatitis tissues.

|  | *Murine AP* | **ADM<sup>1</sup>** | **Proliferative** | **Apoptotic** |
| --- | --- | --- | --- | --- |
| **Cellular markers** |  | Sox9/Amylase | Ki67 | cl.caspase8 |
| **% of ST6GAL1-positive cells<sup>2</sup>** |  | 71.10 ± 15.52 | 31.19 ± 18.26 | 0.00 ± 0.00 |

|  | *Human CP* | **ADM<sup>1</sup>** | **Proliferative** | **Apoptotic** |
| --- | --- | --- | --- | --- |
| **Cellular markers** |  | Sox9/Amylase | Ki67 | TUNEL |
| **% of ST6GAL1-positive cells<sup>2</sup>** |  | 78.48 ± 15.23 | 29.00 ± 14.26 | 3.26 ± 4.63 |

<sup>1</sup> ADM-like cells are marked by dual expression of Sox9 and Amylase.
<sup>2</sup> The % of ST6GAL1-positive cells was calculated as the proportion of ST6GAL1-expressing cells with co-expression of the indicated cellular markers as compared with all ST6GAL1-expressing cells.
*Murine AP*: pancreatitis was induced in mice (n = 5) via cerulein injections and pancreatic tissue was stained for ST6GAL1, Sox9, amylase, Ki67, and cl. caspase8.
*Human CP*: a human pancreatitis tissue array (n = 59) was stained for ST6GAL1, Sox9, amylase, Ki67. A TUNEL assay was utilized to determine acinar cell apoptosis in human samples.

### ST6GAL1 promotes the activation of EGFR, ERK and AKT and upregulation of anti-apoptotic proteins

EGFR signaling induces Sox9 expression and ADM (9–12, 51), and protects against cerulein-induced pancreatitis (52, 53). We thus examined EGFR activation (p-EGFR) in the acini of healthy murine pancreas (saline-injected) or AP tissues (cerulein-injected). We compared WT mice with SC mice that have transgenic expression of ST6GAL1 in the pancreas (*Pdx1-Cre; LSL-ST6GAL1*). In WT mice (littermate controls expressing *LSL-ST6GAL1*), healthy acinar cells lacked detectable p-EGFR whereas robust activation of EGFR was observed in acinar cells from the cerulein cohort (Figure 3A). These data are in line with other studies demonstrating EGFR activation in the ADM-like cells within pancreatitis tissues (3, 4, 54). Of note, WT cells with activated EGFR in AP tissues co-expressed endogenous ST6GAL1. SC acinar cells in AP tissues likewise displayed EGFR activation, as expected, however EGFR was also strongly activated in the healthy SC acinar cells of saline controls, in stark contrast to healthy WT acinar cells (Figure 3A). The levels of total EGFR were not affected by ST6GAL1 transgene expression.

**Figure 3.**
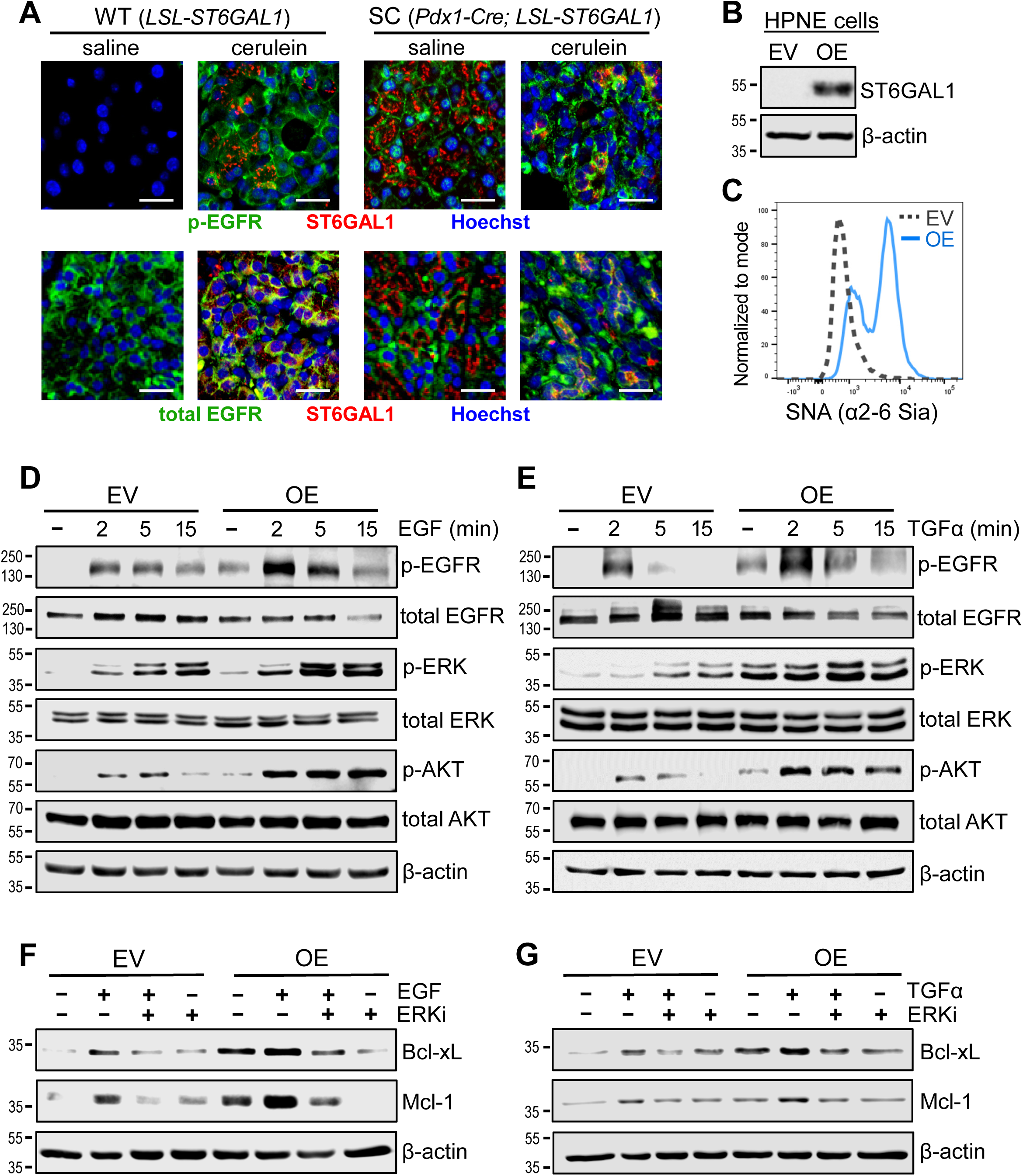
ST6GAL1 promotes the basal and ligand-dependent activation of EGFR, ERK and AKT, and enhances the expression of anti-apoptotic proteins. **(A)** SC mice (*Pdx1-Cre*; *LSL-ST6GAL1*) and WT littermate controls (*LSL-ST6GAL1*) were injected with saline or cerulein to induce AP. Upper panels: IF staining for phospho-EGFR (p-EGFR, green) and ST6GAL1 (red). Lower panels: IF staining for total-EGFR (green) and ST6GAL1 (red). p-EGFR was detected in the cerulein-treated WT and SC groups, as expected, however p-EGFR was also detected in the healthy pancreas of SC, but not WT, mice. Scale bars = 25 μm; original magnification, 200×. **(B)** HPNE cells with undetectable endogenous ST6GAL1 were stably transduced with lentivirus encoding ST6GAL1 to overexpress the enzyme (OE), or with an empty vector (EV) lentiviral construct. **(C)** OE cells displayed increased surface α2-6 sialylation relative to EV cells, as measured by SNA staining and flow cytometry. **(D-E)** Cells were treated with 50 ng/ml EGF (**D**) or TGFα (**E**) for varying time points. Cell lysates were immunoblotted for p-EGFR, total-EGFR, p-ERK1/2, total-ERK1/2, p-AKT, and total-AKT. **(F-G)** Cells were treated with 50 ng/ml EGF (**F**) or TGFα (**G**) for 24 hours. For some experiments, an ERK1/2 inhibitor (ERKi, SCH772984) was included. Treated cells were immunoblotted for the Bcl-2 family anti-apoptotic proteins, Bcl-xL and Mcl-1.

To delve into the molecular mechanisms by which ST6GAL1 reprograms cell phenotype, we used the HPNE nonmalignant pancreatic epithelial cell line (55). ST6GAL1 expression was not detected in parental HPNE cells, consistent with the negligible expression of ST6GAL1 in normal pancreatic epithelium. We therefore generated a cell model with stable overexpression of ST6GAL1 (OE), or a control line with an empty vector (EV) construct (Figure 3B). Increased ST6GAL1-mediated α2-6 sialylation of OE cells was verified by staining cells with *Sambucus Nigra Agglutinin* (SNA), a lectin specific for α2-6-linked sialic acids (Figure 3C). The EV and OE HPNE cell lines were monitored for the activation of EGFR following treatment with EGF or TGFα. OE cells displayed greater activation of EGFR in response to both EGF (Figure 3D) and TGFα (Figure 3E). Intriguingly, basal (ligand-independent) activation of EGFR was also elevated in OE cells, which aligns with our prior studies (21, 24, 56). Along with heightened EGFR activation, OE cells exhibited increased basal and ligand-induced activation of ERK and AKT (Figure 3D and E). Furthermore, OE cells had increased levels of the anti-apoptotic proteins Mcl-1and Bcl-xL, both basally and in response to EGF and TGFα (Figures 3F and G). The expression of Mcl-1 and Bcl-xL was attenuated in cells pre-incubated with an ERK inhibitor (Figures 3F and G). The results in Figure 3 suggest that ST6GAL1 is a strong activator of survival pathways associated with EGFR signaling.

### ST6GAL1 activity enhances the survival of cells exposed to stress

Our prior studies of cancer cells implicated ST6GAL1 in protection against diverse cytotoxic stressors. We hypothesized that this function would be conserved in non-malignant epithelial cells. We therefore monitored the survival and proliferation of HPNE cells exposed to a variety of stress-inducing stimuli. We first measured the proliferation of cells grown in normal growth media containing 5% FBS (“serum-high”), or serum-depleted media containing 0.5% FBS (“serum-low”). In serum-high media, a greater number of OE cells was apparent by the 5-day time point (Fig 4A). A more dramatic difference in proliferation was observed under serum-low conditions, with higher numbers of OE vs. EV cells at all time points tested after initial seeding (Fig 4A). Cell response to serum deprivation was also evaluated by quantifying the number of cells in S-phase at 24 and 72 hours. In serum-low conditions, significantly more S-phase cells were detected in OE cells, whereas there was no appreciable difference when serum was replete (Figure 4B, scatter plots in upper panels, bar graphs in lower panels). We then examined the growth of cells seeded at a very low density that precluded the initial formation of cell-cell junctions, similar to conditions for a clonogenics assay. However, because HPNE cells do not growth as discrete colonies, we quantified total cell number rather than colony number. After 3 weeks of growth in low-density conditions, a greater number of OE cells compared to EV cells was detected in cells grown in low serum (Figure 4C). There was a trend toward increased OE cell growth in serum-high media, but results were not significant. Finally, soft agar assays were employed to show that OE cells exhibited better anchorage-independent growth than EV cells (Figure 4D). The results in Figure 4 suggest that ST6GAL1 promotes the survival and growth of cells exposed to stress.

**Figure 4.**
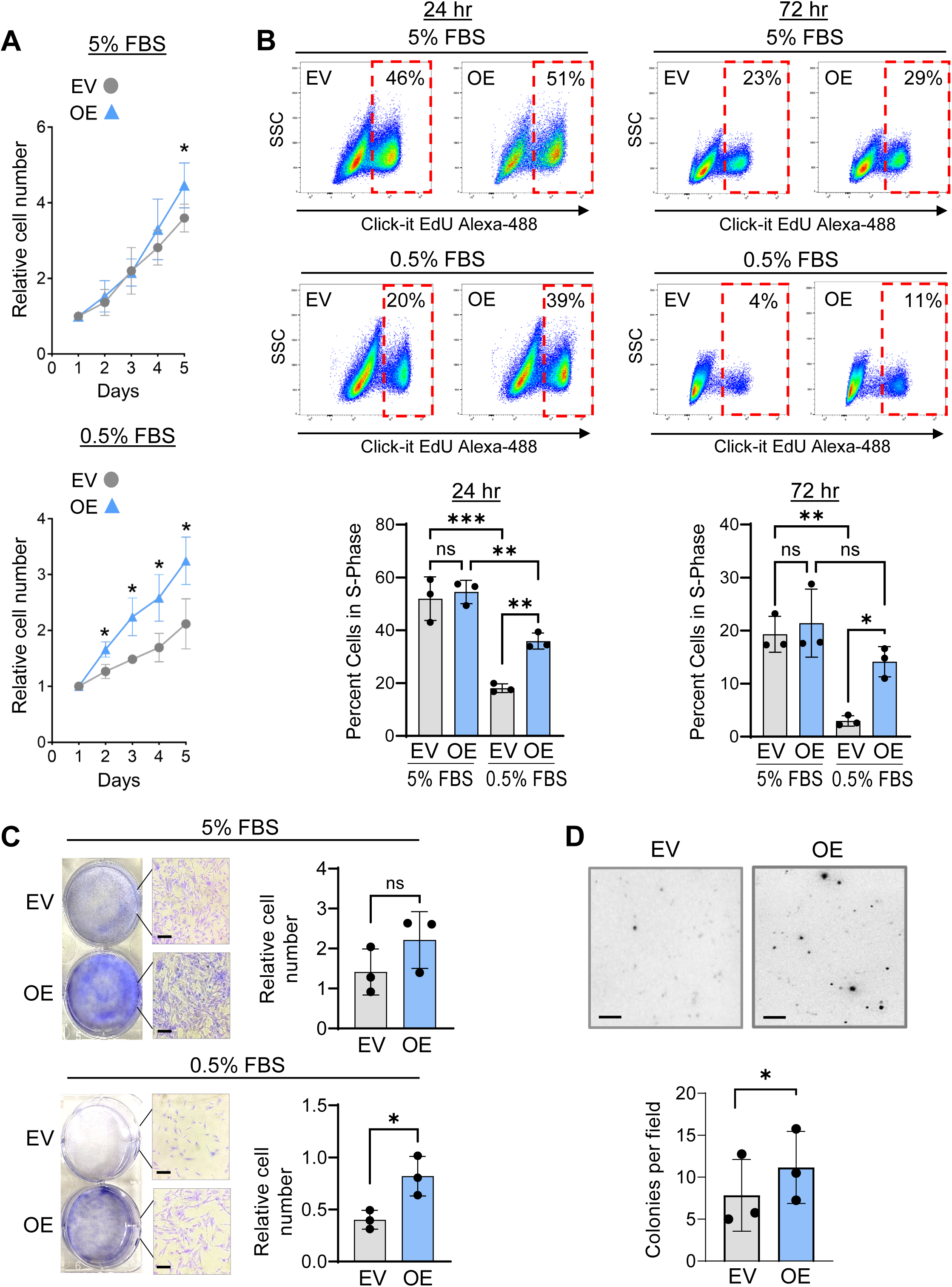
ST6GAL1 enhances the survival of cells exposed to stress. **(A)** HPNE EV and OE cells were monitored for proliferation in media containing either 5% FBS (normal serum level, upper panel) or 0.5% FBS (low serum conditions, lower panel). The total number of cells was quantified using a Cyquant assay. **(B)** HPNE cells were grown in 5% FBS or 0.5% FBS for 24 or 72 hours. Click-it Edu staining was used to label cells in S phase. The percentage of cells in S-phase is indicated in the red dashed boxes. SSC = side scatter. **(C)** HPNE cells were cultured at very low seeding densities in 5% or 0.5% FBS and allowed to grow for 3 weeks. Cells were stained with crystal violet, and relative cell number was measured by solubilizing cells in 10% acetic acid and reading absorbance at 590 nm. Scale bars = 25 μm; original magnification, 100×. **(D)** HPNE cells were subjected to soft agar assays. Anchorage-independent growth was quantified by counting the number of colonies. Scale bars = 500 μm; original magnification, 40×. For all data in Figure 4, three independent experiments were conducted. Results are depicted as mean ± S.D., with a Student’s t-test or 2-way ANOVA employed for statistical analysis. \**p* < 0.05. \*\**p* < 0.005. \*\*\**p* < 0.0005.

### ST6GAL1 activates Receptor Tyrosine Kinase signaling networks

The activation of RTKs is often associated with a pro-growth cell phenotype, and constitutive activation of select RTKs is prevalent during malignant transformation (57). Having shown ST6GAL1 activates EGFR, we screened more broadly for kinase activity by performing a kinomics assay. Kinomics assays provide an unbiased assessment of the activation of numerous cytosolic and membrane-associated tyrosine and serine/threonine kinases. A heatmap of the kinomics data from HPNE cells shows that many kinases were differentially activated in the EV and OE lines (Figure 5A). In particular, OE cells displayed greater kinase activation in a region of the heatmap (dashed box) that includes read-outs for multiple oncogenic RTKs including EGFR, MET, PDGFRβ, ERBB2, and AXL. Along with EGFR, the MET, PDGFRβ and ERBB2 receptors are known to be activated by ST6GAL1-mediated sialylation (43, 58, 59). Figure 5B indicates the relative level of tyrosine kinase activity. Among the > 40 tyrosine kinases that differed between EV and OE cells, all displayed greater activation in the OE line. The RTKs activated in OE cells included EGFR, PDGFRβ, PDGFRα, FGFR3/4, MET, ERBB2, RON, and MER. Statistical values for the top 10 tyrosine kinases with elevated activity in OE cells are shown in Figure 5C. In contrast to tyrosine kinases, serine/threonine kinases were generally less active in OE cells (Figure S1). Serine/threonine kinases with reduced activity in OE cells included RSK2, ROCK1 and 2, PKCα, and PKG1 and 2 (Figure S1).

**Figure 5.**
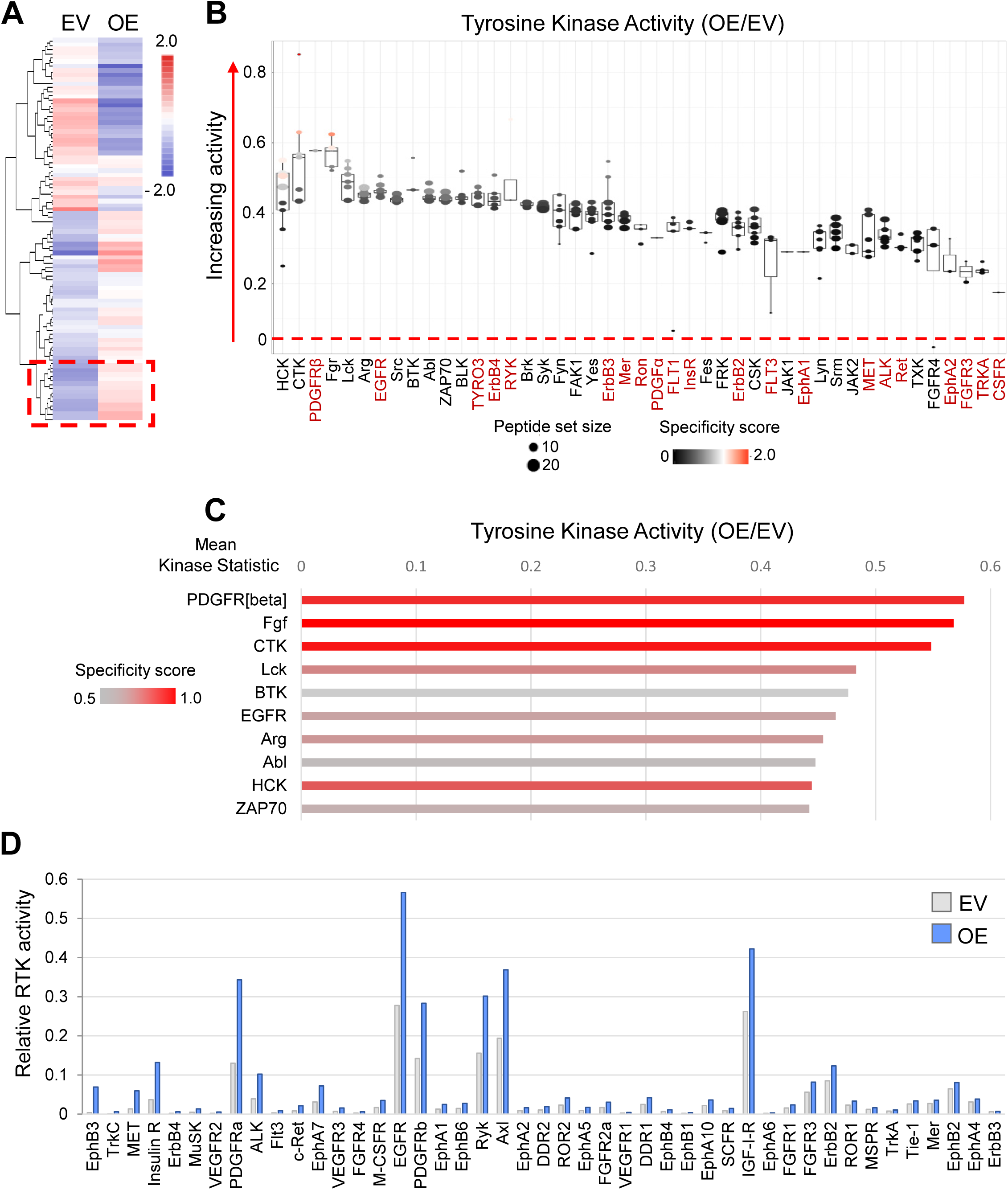
ST6GAL1 promotes the activation of RTKs including EGFR and other growth factor receptors. **(A)** A kinomics assay was used to monitor the activity of membrane-associated and cytosolic tyrosine and serine/threonine kinases. The heatmap shows the kinomic peptide signal of HPNE EV and OE samples. Data displayed are colored by change per-peptide from Log2 mean. Red dashed box indicates a region of the heatmap rich in read-outs for RTKs. **(B)** Kinases identified using whole-chip (all peptide) comparison from the relative level of tyrosine kinase activation in OE vs. EV cells. The Y-axis indicates the amount and direction of kinase change with a positive score indicating an increase in kinase activity in OE cells. RTKs are labelled in red. High specificity score indicates strong activity of the associated kinase on its intended target peptides. **(C)** Statistical values for the top 10 tyrosine kinases elevated in OE vs. EV cells. The mean kinase statistic indicates the overall change in activity for a particular kinase. **(D)** An RTK array was used to measure RTK activity in HPNE EV and OE cells.

To validate the kinomics data, we probed for the activity of RTKs using an RTK array. Figure 5D shows that the great majority of RTKs on the array had greater activation in OE vs. EV cells. RTKs activated in OE cells included EGFR, PDGFRβ, FGFR4, VEGFR2 and 3, MET, ERBB4, Insulin receptor, TrkC, and multiple Eph receptors. The results in Figure 5 suggest that ST6GAL1 promotes the activation of many growth factor receptors and other RTKs that are associated with cell survival and proliferation.

### ST6GAL1 inhibits apoptosis induced by the Fas and TNFR1 death receptors

In conjunction with the activation of growth factor receptors, acinar cells within the pancreatitis milieu must resist cell death induced by inflammatory cytokines such as the death receptor ligands, TNFα and FasL. These ligands bind to TNFR1 and Fas, respectively, to induce acinar cell death through both apoptosis and necroptosis (30). Interestingly, acinar cells undergoing ADM are known to be resistant to Fas-mediated apoptosis (60). To assess the role of ST6GAL1 in death receptor signaling, HPNE EV and OE cells were treated with TNFα or FasL and monitored for the activation of the initiator caspase, caspase 8 and the executioner caspase, caspase 3. Activation of caspase 8 is a hallmark of the extrinsic, death receptor-mediated apoptotic pathway. OE cells had sharply reduced caspase activation in response to TNFα (Figure 6A) and FasL (Figure 6B). Lactate dehydrogenase (LDH) release assays similarly showed that ST6GAL1 protected against cell death induced by TNFα and FasL (Figure 6C and D).

**Figure 6.**
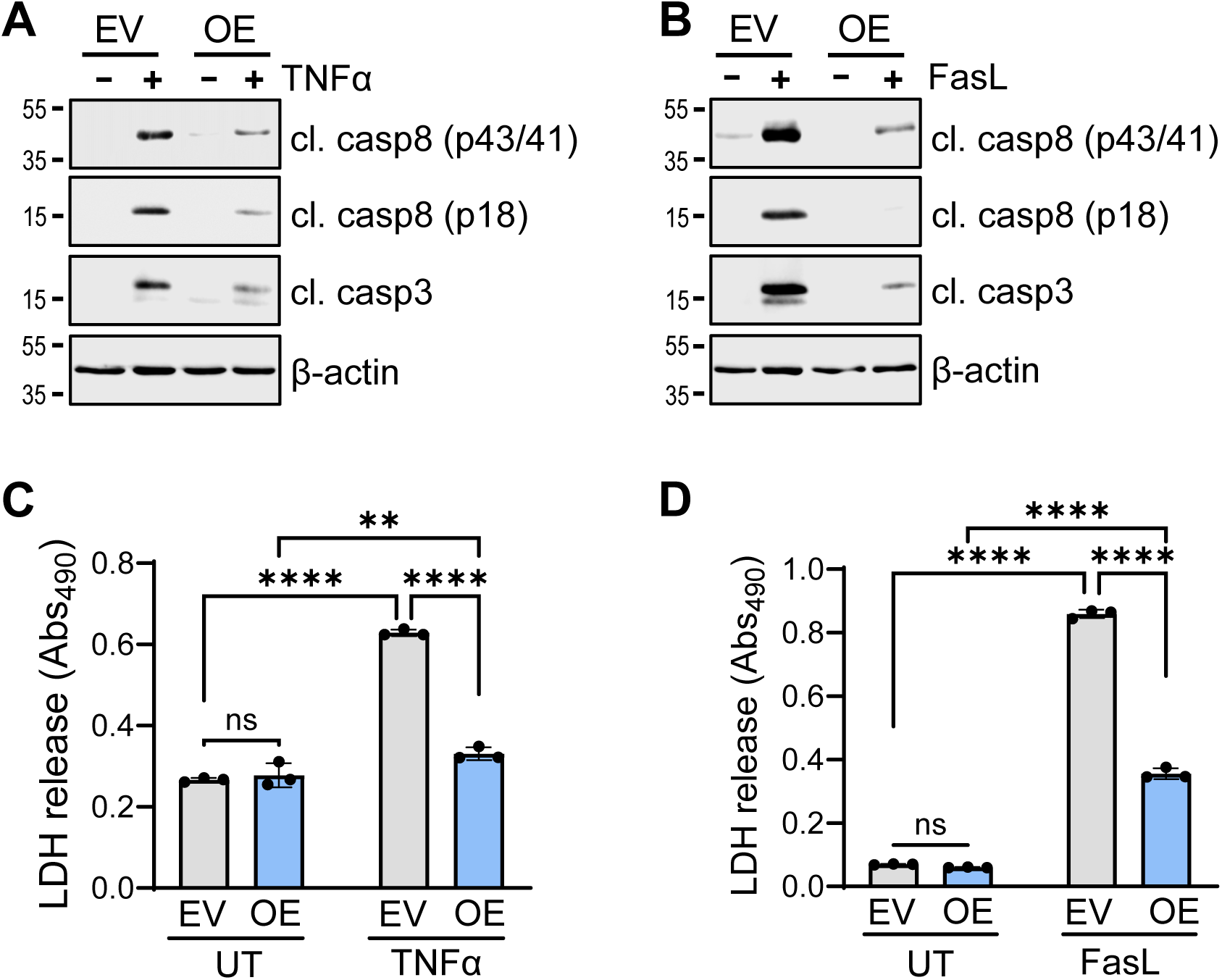
ST6GAL1 inhibits TNFα and FasL dependent death receptor-mediated apoptosis. (**A-B**) HPNE cells were treated for 24 hours with 100 ng/mL TNFα **(A)** or 10 ng/mL FasL (**B**). Cell lysates were immunoblotted for cl. casp8 (p43/41 and p18 cleavage products) and cleaved caspase 3 (cl. casp3). **(C-D)** Conditioned media was collected 24 hours after treating cells with TNFα (**C**) or FasL (**D**). Enzyme-coupled assays were used to quantify LDH levels in the media. LDH release graphs represent mean ± S.D. from 3 independent experiments, data analyzed by 2-way ANOVA. \*\**p* < 0.01. \*\*\*\**p* < 0.0001.

### Pro-survival signaling networks are activated in organoids derived from ST6GAL1 transgenic mice

To evaluate survival signaling in another epithelial model, we used organoids derived from the pancreata of WT and SC mice. Our prior studies revealed that, as compared with WT organoids, SC organoids have an ADM-like phenotype as indicated by: (1) drastically reduced expression of mature acinar genes; (2) increased expression of stem and ductal genes; and (3) more robust organoid-forming capability and organoid growth (32). WT and SC organoids were treated with EGF or TGFα, or left untreated (basal activation), and monitored for EGFR activation by IF microscopy. SC organoids demonstrated enhanced basal and ligand-induced EGFR activation relative to WT organoids (representative images in Figure 7A, quantification in Figure 7B). Congruently, SC organoids had increased basal and ligand-induced activation of ERK (Figure 7C and D) and AKT (Figure 7E and F). SC organoids also displayed an enrichment in the expression of Mcl-1 and Bcl-xL (Figure 7G and H). We then examined death receptor-induced apoptosis. Following treatment with TNFα or FasL, SC organoids had decreased activation of caspases 8 and 3 (Figures 7I and J), and reduced release of LDH (Figures 7K and L). These data indicate that ST6GAL1 activates pro-survival signaling cascades and simultaneously inhibits death receptor-mediated apoptosis in primary pancreatic epithelial cells.

**Figure 7.**
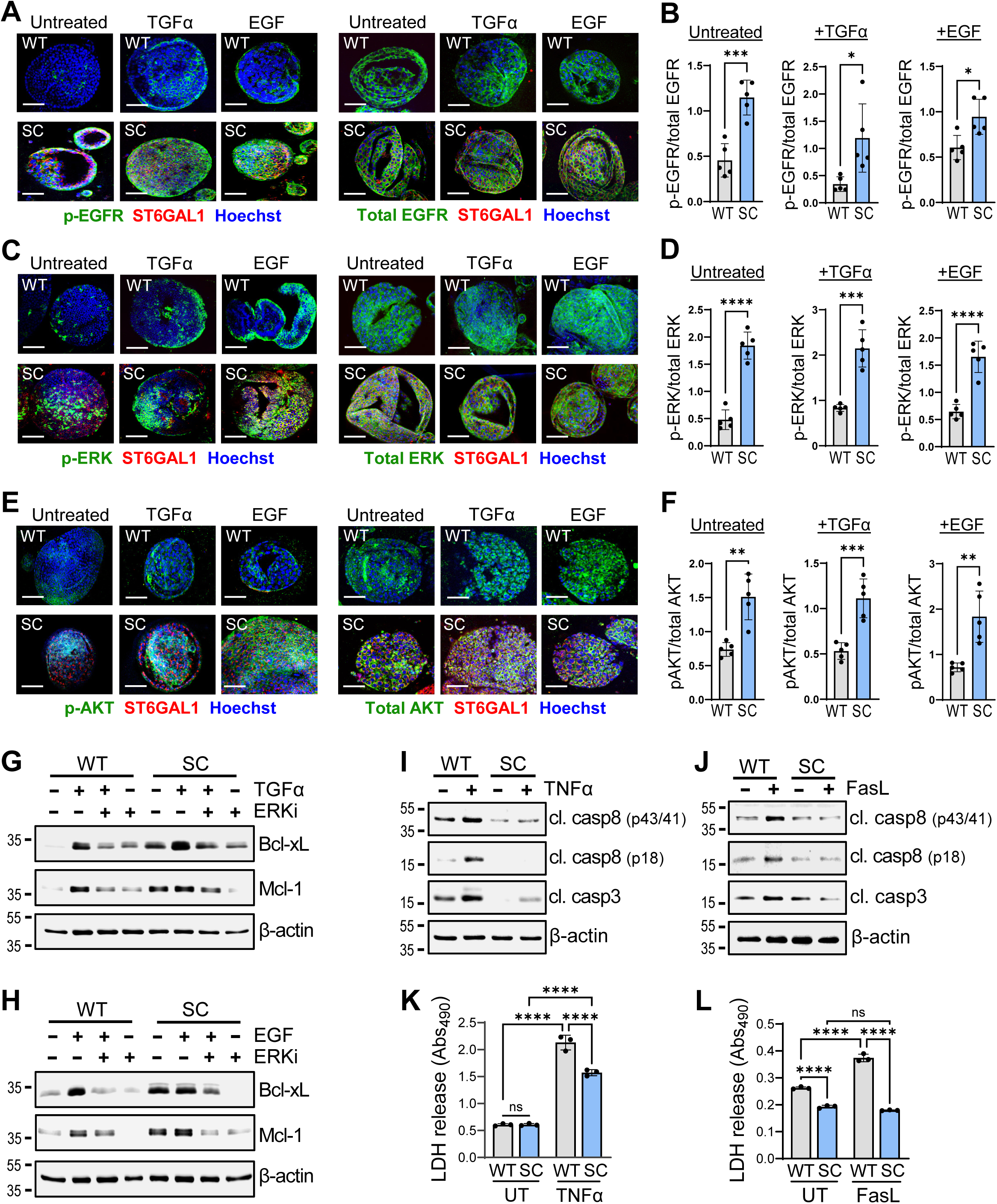
Greater activation of pro-survival signaling networks in pancreatic organoids generated from SC vs. WT mice. **(A)** Representative images of WT (top) and SC (bottom) organoids either left untreated or treated with 100 ng/mL TGFα or EGF. Left 6 panels depict organoids stained for p-EGFR (green) and ST6GAL1 (red), whereas right 6 panels show organoids stained for total EGFR (green) and ST6GAL1 (red). **(B)** Quantification of EGFR activation. The fluorescence intensity of p-EGFR was normalized to the intensity of Hoechst. The intensity of total EGFR was likewise normalized to Hoechst. The normalized p-EGFR value was then divided by the normalized total EGFR value to obtain a measure of EGFR activation. Graphs depict the p-EGFR/total EGFR ratios for untreated (basal activation), TGFα-treated and EGF-treated organoids. (**C**) Representative images of untreated, TGFα-treated and EGF-treated WT and SC organoids stained for p-ERK or total ERK (green) and ST6GAL1 (red). (**D**) The activation of ERK was quantified by calculating p-ERK/total ERK ratios as described for EGFR in panel B. (**E**) Representative images of WT and SC organoids stained for p-AKT or total AKT (green) and ST6GAL1 (red). (**F**) The activation of AKT was quantified by calculating p-AKT/total AKT ratios, as described in panel B. Scale bars in panels **A**,**C**,**E** = 100 μm; original magnification, 200×. For panels **B**, **D**, **F**, quantification was performed on 5 randomly selected organoids. At least two independent experiments were performed for each signaling molecule. Graphs depict mean ± S.D., with data analyzed using a Student’s t-test. **(G-H)** Organoids were treated with 100 ng/mL TGFα (**G**) or EGF (**H**) for 24 hours in the presence or absence of an ERK1/2 inhibitor (ERKi, SCH772984). Organoid homogenates were immunoblotted for the anti-apoptotic proteins Bcl-xL and Mcl-1. **(I-J)** Organoids were treated for 24 hours with 100 ng/mL TNFα **(I)** or 10 ng/mL FasL (**J**) and immunoblotted for cl. casp8 (p43/41 and p18 cleavage products) and cl. casp3. **(K-L)** Media was collected from organoids treated for 24 hours with TNFα (**K**) or FasL (**L)** and analyzed for LDH release. LDH release graphs represent mean ± S.D. for 3 independent experiments, with data analyzed by 2-way ANOVA. \**p* < 0.05. \*\**p* < 0.005. \*\*\**p* < 0.0005. \*\*\*\**p* < 0.0001.

### ST6GAL1 promotes pancreatic tissue recovery following acute pancreatitis

Results from the HPNE and organoid models suggested that the upregulation of ST6GAL1 in acinar cells may foster a pro-survival phenotype important for regenerating the exocrine pancreas. To test this hypothesis, we monitored tissue recovery in WT and SC mice treated with L-arg to induce AP. We postulated that the acinar cells of SC mice would be primed for a reparative response given their more ADM-like phenotype (32). WT and SC mice were injected with L-arg and then pancreata were harvested after 4, 6 or 10 days (D4, D6, or D10) (Figure 8A). Pancreatic tissues were stained by H&E to assess histomorphological characteristics such as acini architecture, inflammation, edema, fibrosis, and necrosis. At 4 days following L-arg injection, substantial damage to the exocrine pancreas was observed in both WT and SC pancreata (Figure 8B). However, at 6 and 10 days, there appeared to be less damage in SC compared with WT mice, consistent with more rapid tissue recovery (Figure 8B). SC pancreata were fully healed by 10 days, whereas WT pancreata, although mostly healed, still retained some regions of damage. We confirmed that endogenous ST6GAL1 levels were increased in the WT L-arg-treated samples in the damaged areas of the pancreata at Days 4, 6, and 10 (Figure 8C). As expected, ST6GAL1 was expressed throughout the pancreata of all SC samples (Figure 8C). To confirm that ST6GAL1 upregulation resulted in enhanced α2-6-linked sialyation of acinar cells, we co-stained tissues with the SiaFind α2,6-specific reagent and for amylase. In the WT L-arg-treated samples, α2-6-sialylation was observed in the ADM-like acinar cells, as identified by their ductal-like morphology with secreted amylase in the lumen of these structures (Figure 8D, arrows). α2-6-sialylation was not detected in areas in which the pancreatic tissue had recovered and demonstrated normal acinar morphology (Figure 8D, asterisks). SC samples exhibited α2-6-sialylation of acinar cells in both the ADM-like and normal acini (Figure 8D). These results suggest that ST6GAL1 is upregulated selectively in damaged acinar cells undergoing ADM. However, once ADM is resolved, the acini revert to their normal, quiescent phenotype and ST6GAL1 is no longer expressed, indicating that the upregulation of ST6GAL1 during AP is transient.

**Figure 8.**
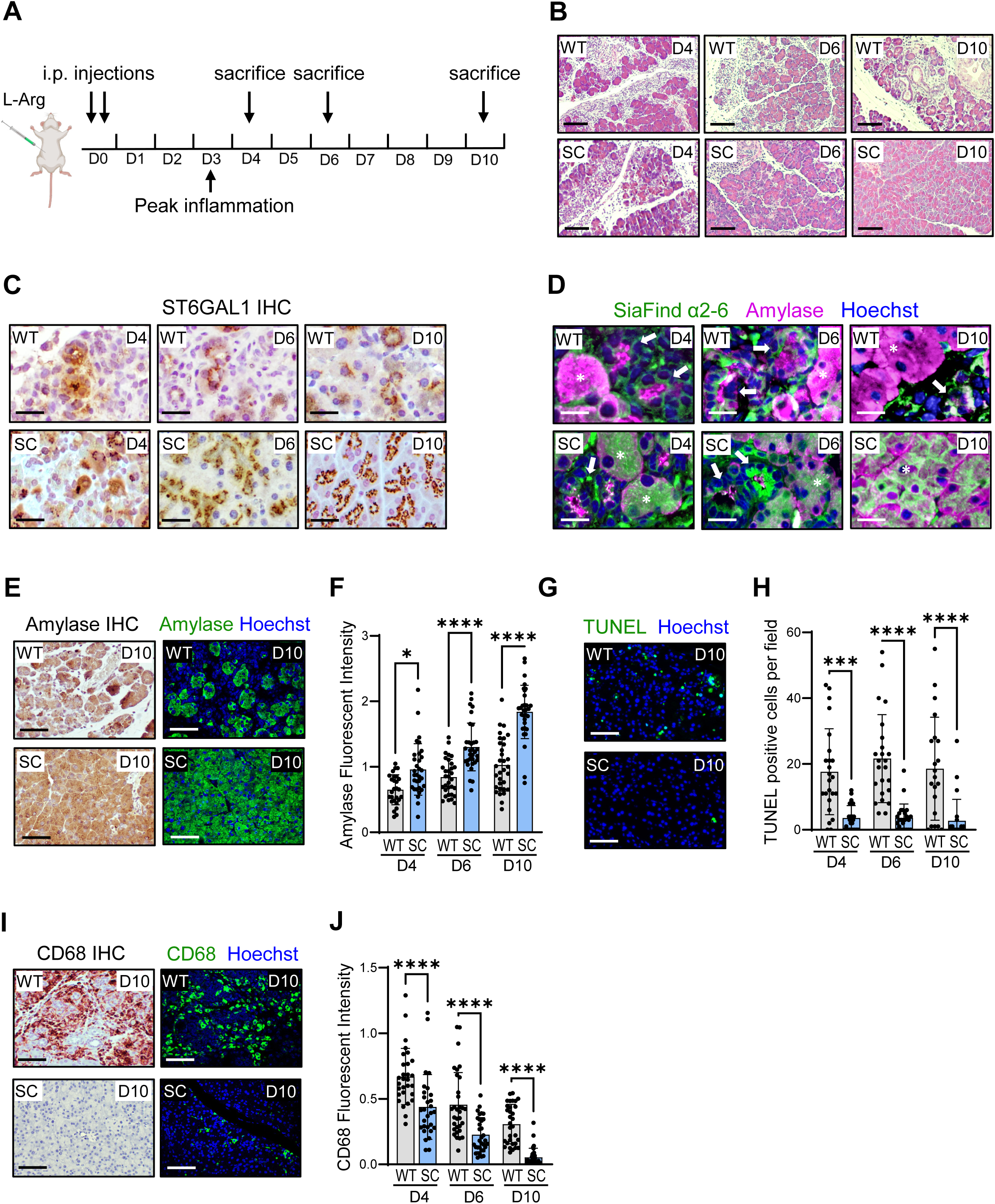
ST6GAL1 promotes pancreatic tissue recovery following induction of AP. **(A)** Schematic showing recovery interval. WT and SC mice were given two i.p. injections of L-arg on Day 0 (D0). Pancreata were harvested for analysis on Days 4, 6 and 10 (D4, D6, D10). **(B)** H&E images of WT (top) and SC (bottom) pancreata at various intervals following L-arg injection. Tissue damage was comparable in WT and SC mice at D4, whereas greater tissue repair was apparent in SC mice at D6 and D10. Scale bars = 100 μm; original magnification, 100×. **(C)** IHC staining showing upregulation of endogenous ST6GAL1 in WT pancreata at D4, D6, and D10 in areas of damaged pancreata (top panels). Constitutive ST6GAL1 expression is noted in SC pancreata, reflecting expression of the ST6GAL1 transgene (bottom panels). Scale bars = 50 μm; original magnification, 200×. **(D)** IF staining of WT tissues (top panels) with the SiaFind α2,6 reagent confirmed upregulation of α2-6-linked sialic acid in ADM-like structures, as indicated by a ductal-like morphology with amylase secretion into the lumen (arrows). Normal-appearing acinar cells (asterisks) lacked α2-6 sialylation. In SC tissues (bottom panels), high α2-6-sialylation was detected in both ADM-like structures (arrows) and normal acinar cells (asterisks), consistent with constitutive expression of the ST6GAL1 transgene. Scale bars = 25 μm; original magnification, 200×. **(E)** Representative amylase IHC and IF images of WT and SC pancreata at D10. Scale bars = 50 μm; original magnification, 200×. **(F)** Quantification of IF images indicates increased amylase levels in SC pancreata at D4, D6, and D10. **(G)** Representative images from TUNEL analysis of WT and SC pancreata at D10. Scale bars = 50 μm; original magnification, 200×. **(H)** Quantification of TUNEL-positive cells indicates reduced apoptosis in SC pancreata at D4, D6, and D10. **(I)** Representative IHC and IF staining for CD68 in WT and SC pancreata at D10. Scale bars = 50 μm; original magnification, 200×. **(J)** Quantification of CD68 expression suggests reduced monocyte/macrophage infiltration in SC pancreata at D4, D6, and D10. Quantification in panels **F**, **H**, and **J** was conducted using 8 mice per group, with 4 (**F** and **H**) or 3 (**J**) distinct areas analyzed per pancreas. Data are depicted as mean ± S.D., and 2-way ANOVAs were used for statistical analyses. \**p* < 0.05, \*\*\**p* < 0.001, \*\*\*\**p* < 0.0001.

Tissues were next stained for amylase as a marker of acinar integrity. At the 10-day time point, WT pancreata maintained areas of acinar disruption, unlike SC pancreata for which the acini had a uniformly normal-appearing morphology (Figure 8E). Additionally, there was quantitatively less amylase staining in WT pancreata at 4, 6 and 10 days, consistent with tissue damage (Figure 8F). We then performed TUNEL staining, which revealed significantly greater acinar cell death in WT vs. SC pancreata at Days 4, 6, and 10 (Figure 8G and H). Finally, staining for CD68 indicated that WT pancreata had increased numbers of monocytes and macrophages compared with SC pancreata at all time points, reflecting increased inflammation (Figure 8I and J). The combined results in Figure 8 suggest that ST6GAL1 promotes more rapid recovery of the exocrine pancreas following pancreatitis-induced tissue damage.

## Discussion

Pancreatitis is associated with significant morbidity, and in severe cases, mortality (1, 2). Most clinical interventions treat disease symptoms but cannot reverse tissue damage. Understanding the molecular processes that underlie pancreatic injury and repair is crucial for the design of more effective treatments. While regenerative processes within the exocrine pancreas have been previously studied (3), the potential contribution of cellular glycosylation has been largely overlooked. Intriguingly, certain glycan structures, including select sialoglycans, are elevated in response to pancreatic inflammation. The sialyl Lewis A tetrasaccharide (sLeA, CA19-9) is increased in AP and CP patient tissues (61, 62), and mice engineered to express sLeA develop AP, which then progresses to CP (63). sLeA is also overexpressed in PDAC and serves as the primary clinical biomarker for disease recurrence and response to treatment (64). Other prominent glycans increased in pancreatitis and PDAC include the Tn and sialyl Tn (sTn) antigens. Mice that express Tn and sTn due to a genetic deletion in the Cosmc chaperone protein (CT1GALT1) develop CP (65), and also exhibit enhanced PDAC progression when crossed to KC mice (66). These findings implicate glycans in both the development of pancreatitis and the transition from pancreatitis to malignancy.

In the present investigation, we highlight another sialoglycan enriched in pancreatitis, the α2-6 sialyllactosamine structure generated by ST6GAL1 (Neu5Acα2,6Galβ1-4GlcNAc). To study ST6GAL1’s function in the pancreas, we previously generated SC mice, which have transgenic expression of ST6GAL and elevated α2-6 sialylation of pancreatic acinar cells (32). In contrast to mice expressing sLeA and Tn/sTn, SC mice do not spontaneously develop pancreatitis. Instead, we find that ST6GAL1 facilitates tissue healing. As reported (32), the acinar cells of SC mice have a ductal, progenitor-like phenotype; we therefore postulated that these cells would be primed to mount a regenerative response upon pancreatic injury. Results herein show that following the induction of AP by L-arg, SC mice displayed accelerated pancreatic tissue recovery when compared with WT mice, evidenced by more rapid restoration of the acini, markedly reduced acinar cell death, and lower immune cell infiltration. These data point to a new function for ST6GAL1 in tissue repair, an area of research for which there is a striking dearth of literature. In one publication by Punch et al. (67), it was shown that ST6GAL1 was necessary for regeneration of the gastrointestinal tract following whole-body irradiation of mice. However, this study utilized a global ST6GAL1 knock-out mouse model, and the mechanism by which ST6GAL1 contributed to tissue recovery was not investigated. Here, we propose a pro-regenerative role for ST6GAL1 that enables restoration of the exocrine compartment following a bout of pancreatitis.

The pancreatitis-induced upregulation of ST6GAL1 was conserved across human AP and CP, and two murine models of AP. Notably, ST6GAL1 was selectively expressed in the ADM-like cells (co-expressing Sox9 and amylase) within pancreatitis tissues. The mechanism responsible for elevated ST6GAL1 expression during pancreatitis may involve signaling by the pro-inflammatory cytokines, IL-1β and IL-6, which are abundant within the inflammatory milieu (68). We previously reported that IL-1β and IL-6, acting through their respective transcription factors, NFκB and STAT3, induce the transcriptional upregulation of ST6GAL1 in both malignant and non-malignant pancreatic cells (e.g., HPNE cells) (69). The IL6/STAT3 pathway plays a particularly prominent part in ADM and PDAC initiation. During pancreatitis, IL-6 is secreted by infiltrating macrophages, damaged acinar cells and several other cell types (70–74). IL-6 then drives the conversion of mature, quiescent pancreatic acinar cells into progenitor-like, ductal cells that can proliferate in response to the IL-6/JAK/STAT3 signaling cascade (75–77). Activation of STAT3 by IL-6 also promotes cell survival and resistance to apoptosis during pancreatitis (78). Indeed, IL-6 signaling stimulates the expression of several anti-apoptotic genes (e.g. Bcl-xL), while suppressing pro-apoptotic genes (77, 79). However, in the presence of oncogenic KRas, IL-6 signaling facilitates tumor initiation. For example, IL-6 is required for the maintenance of pancreatic cancer precursor lesions (<u>Pan</u>creatic Intraepithelial <u>N</u>eoplasms, PanINs) in the presence of inflammation, and deletion of IL-6 decreases the proliferative capacity of tumor and stromal cells (77).

In cancer cells, ST6GAL1 is a well-known driver of cell survival, conferring protection against hypoxia (80), serum withdrawal (81) and chemotherapeutics (38, 82). However, very little is known about ST6GAL1’s function in normal epithelium. One mechanism by which ST6GAL1 may aid in pancreatic tissue repair is through preventing acinar apoptosis, thereby increasing the pool of cells available for restoration of the acini. Our analyses of human and murine pancreatitis tissues revealed that ST6GAL1-expressing cells lacked expression of apoptotic markers, whereas many of these cells expressed Ki67, indicating re-entry into the cell cycle. Furthermore, SC mice exposed to L-arg-induced AP had significantly fewer apoptotic acinar cells when compared with WT mice. We posited that apoptosis-resistance was mediated, at least in part, by the α2-6 sialylation of the TNFR1 and Fas death receptors. In both HPNE cells with modulated ST6GAL1 expression and organoids from WT and SC pancreata, high ST6GAL1 expression was found to inhibit caspase activation and cell death induced by TNFα and FasL. The TNFα-TNFR1 axis is a major contributor to pancreatitis-associated pathogenesis (30), and therapeutic interventions that block TNFα activity reduce the severity of AP in animal models (83, 84).

As another key survival mechanism, ST6GAL1 promotes the activation of EGFR, which is one of the main drivers of acinar de-differentiation, survival and proliferation during ADM (9–12). We found that EGFR was activated in the acinar cells of SC mice (i.e., in the absence of pancreatitis), indicating that ectopic ST6GAL1 expression was sufficient to induce EGFR activation. Moreover, organoids and HPNE cells with high ST6GAL1 expression had increased basal and ligand (EGF, TGFα)-stimulated activation of EGFR and its downstream mediators, AKT and ERK. ST6GAL1-expressing cells also displayed an ERK-dependent enrichment in the expression of the Mcl-1 and Bcl-xL anti-apoptotic proteins, an event which may synergize with ST6GAL1’s inhibitory effects on the TNFR1 and Fas death receptors. Phenotypically, ST6GAL1 enhanced the activity of many growth-associated RTKs (PDGFRβ, MET, ERBB2, etc.), and protected cells from various stress-induced stimuli including serum growth factor deficiency, low cell density and anchorage-independent cell growth. These collective findings suggest that ST6GAL1 imparts a pro-survival, pro-growth cellular program that may be critical for tissue repair.

While the current study focused on ST6GAL1’s role in epithelial reprogramming, the potential involvement of ST6GAL1 in immune resolution following pancreatitis may be a fruitful future direction. ST6GAL1 was first identified as an acute phase response protein more than 40 years ago (85), and its role in immune tolerance has long been appreciated. In part, this is due to the ST6GAL1-dependent generation of sialoglycan ligands for the Siglec family of immune checkpoint receptors. Siglec expression is largely restricted to immune cells and, like other checkpoint proteins (e.g. PD1), inhibitory Siglecs have ITIM motifs that function to dampen immune cell signaling (86, 87). ST6GAL1 is the principal enzyme that generates α2-6 sialylated *N*-glycans recognized by Siglec 2 (CD22) on B lymphocytes, an interaction that suppresses B cell activation (88). Siglec 2 is also expressed on certain subsets of macrophages (89) and dendritic cells (DCs) (90, 91). Additionally, Siglec 3 (CD33) and Siglec 10 have been reported to recognize α2-6-linked sialoglycans (92, 93), although these Siglecs engage other types of sialylated structures as well (86, 87). Siglec 3 is expressed by monocytes, neutrophils, and DCs, whereas Siglec 10 is expressed by monocytes, macrophages, B cells, and a subset of DCs and natural killer (NK) cells (87). Through the activation of these (and possibly other) Siglecs, ST6GAL1-mediated sialylation has broad potential to rewire the immune landscape in order to restore tissue homeostasis. Further studies are warranted to determine whether ST6GAL1 mediates healing of the pancreas through its coordinate regulation of acinar and immune cells.

In summary, results from this investigation suggest that the pancreatitis-induced upregulation of ST6GAL1 functions to promote tissue recovery by enhancing acinar cell survival. ST6GAL1’s role in epithelial biology has primarily been studied in cancer; the concept that ST6GAL1 is important for tissue regeneration breaks new ground. The findings in this report offer novel insights into the mechanisms by which α2-6 sialylation remodels epithelial cell phenotype to effectuate a reparative response to organ injury.

## Methods

### ST6GAL1 IHC on human pancreatic tissues

Pancreatitis tissues from AP and CP patients or adjacent healthy tissues were obtained from either the UAB Tissue Biorepository or US Biomax Inc. (tissue microarrays, BBS14011 and PA485). Sections were IHC-stained for ST6GAL1 as reported (38, 45). Briefly, sections were subjected to antigen retrieval using Antigen Unmasking Solution, Citric Acid Based (Vector Laboratories, H-3300); blocked with 2.5% horse serum for 1 hour; and incubated overnight at 4°C with goat polyclonal antibody against ST6GAL1 (R&D Systems, AF5924). Sections were incubated with ImmPRESS-HRP horse anti-goat IgG (Vector Laboratories, MP-7405) for 1 hour and developed using ImmPACT®NovaRED (Vector Laboratories, SK-4805). Images were captured with a Nikon 80i Eclipse microscope and processed with Nikon NIS-Elements imaging software (BR 4.30.02). ST6GAL1-positive cells within the exocrine pancreas were quantified.

### Cell Culture

The hTERT-HPNE (“HPNE”) cell line was purchased from ATCC (CRL-4023). HPNE cells were transduced with lentivirus encoding *ST6GAL1* (Genecopoeia, LPP-M0351-Lv197-100) or a negative control lentivirus (Genecopoeia, LPP-NEG-Lv197-100). Lentiviral transduction was performed using an MOI of 5, and stable polyclonal populations were selected using 10 μg/ml Blasticidin S HCl (Gibco, A1113903). ST6GAL1 expression was confirmed by western blot and also by SNA staining to confirm increased α2-6 sialylation. For the SNA staining protocol (81), cells were blocked with 1% BSA in PBS and then incubated with 20 μg/ml of SNA-FITC (Vector Laboratories, FL-1301–2). SNA-FITC staining was quantified using a LSRII flow cytometer (BD Biosciences).

HPNE cells were cultured in media composed of 75% DMEM without glucose (Sigma, D-5030 with additional 2 mM L-glutamine and 1.5 g/L sodium bicarbonate) and 25% Medium M3 Base (Incell Corp, M300F-500). Complete growth medium included 5% fetal bovine serum (FBS), 10 ng/ml human recombinant EGF (R&D Systems, 236-EG), 5.5 mM D-glucose (1g/L), and 750 ng/ml puromycin. For cell signaling assays (EGFR, AKT, ERK activation), EGF was removed from the growth media several hours before experiments. For the serum-depletion experiments, EGF was excluded from the growth media and the FBS concentration was reduced to 0.5%

### Organoids

Organoids were generated from SC (*Pdx1-Cre*; *LSL-ST6GAL1*) and WT (*LSL-ST6GAL1*) mice according to a standard protocol (94). Briefly, pancreata were minced and placed in collagenase I solution (Gibco) for 45 min with intermittent trituration. Samples were passed through a 70 µM cell strainer (Thermo Fisher Scientific, 22-363-548), and after washing, dispersed cells were resuspended in Matrigel (Corning, 354234). A total of 15 μL of the cell suspension was placed in a 24-well plate. The plate was inverted and the Matrigel allowed to solidify at 37°C for 12 min. Subsequently, cultures were grown for 3 days in organoid media (95), specifically, DMEM containing 50% L-WRN–conditioned media with the following supplements: 10 μM ROCK inhibitor (Selleck Chemicals, S1049), 10 μM TGF-βR1 inhibitor (Selleck Chemicals, SB431542), and 10 mM nicotinamide (LKT Labs, N3310). Organoid cultures were used within 9 passages.

### Induction of pancreatitis in mouse models

#### Cerulein AP

To induce pancreatitis in some experiments, mice were starved overnight and treated with 7 intraperitoneal (i.p.) injections of cerulein (75 μg/kg; R&D Systems, 6264), or saline control, at hourly intervals on 2 alternating days. On day 4, mice were euthanized and pancreata collected for histological analysis.

#### L-arg AP

To induce pancreatitis in pancreatic recovery experiments, WT or SC mice were injected i.p. with 2 doses of 4.5 g/kg L-arg in 0.9% saline (pH 7.0) administered an hour apart. Following L-arg injections, 2 injections of 0.9% saline (pH 7.0) were administered i.p. an hour apart. Mice were monitored and allowed to recover under a heat lamp for 2 hours after final saline injection. Mice were euthanized at 4, 6, or 10 days after injection and pancreata were collected for histological analysis. Eight mice were used for each group. For both experimental models, mice were injected at 2 months of age.

### Histological analysis of pancreatic tissue

#### IHC

Paraffin-embedded tissues were processed for antigen retrieval and IHC-stained for ST6GAL1 (R&D Systems, AF5924), Pancreatic Alpha Amylase (Abcam, ab189341), and CD68 (CST, 97778). Tissue sections were subjected to antigen retrieval with Antigen Unmasking Solution, Citric Acid Based (Vector Laboratories, H-3300) and then blocked with 2.5% horse serum for 1 hour. Sections were incubated with primary antibodies at 4°C overnight, followed by horse anti-goat (Vector Laboratories, MP-7405) or anti-rabbit (Vector Laboratories, MP-7401) ImmPRESS-HRP IgG for 1 hour at room temperature. Reactions were developed using ImmPACT®NovaRED (Vector Laboratories, SK-4805). Sections were counterstained with hematoxylin (Vector Laboratories, H-3404). Slides were mounted using VectaMount® PT Permanent Mounting Medium (Vector Laboratories, H-5000) and images were captured using a Nikon 80i Eclipse microscope.

#### IF staining of pancreatic tissues

Tissue processing and antigen retrieval for paraffin-embedded tissues were achieved as described above. Tissue sections were incubated overnight at 4°C with antibodies against ST6GAL1 (R&D Systems, AF5924), Sox9 (Abcam, ab185230), Pancreatic Alpha Amylase (Abcam, ab189341), Ki67 (Abcam, ab16667), Cleaved Caspase 8 (CST, 8592), EGFR (Abcam, ab52894), and Phospho-EGFR (pTyr-1068, Abcam, ab40815). Primary antibody dilutions were followed as recommended by manufacturer for all antibodies listed except for ST6GAL1 antibody, where a 1:75 dilution was used. Slides were then incubated with species-appropriate Alexa Fluor secondary antibodies (Thermo Fisher Scientific) for 1 hour at 37°C, followed by incubation in Hoechst solution (BD Biosciences, 56190) for 5 min at RT. Slides were mounted using VECTASHIELD Vibrance® Antifade Mounting Medium (Vector Laboratories, H-1700). For some experiments, sequential antibody staining was performed with ST6GAL1 and Sox9 or Ki67 being stained first, followed by fixation with 4% paraformaldehyde in Tris-buffered saline (TBS) for 10 min, and then staining for Pancreatic Alpha Amylase. For TUNEL staining, a Click-iT™ Plus TUNEL Assay (Thermo Fisher Scientific, C10617) was used. Briefly, tissues were incubated with TdT reaction mixture for 60 min at 37°C and then the Click-iT™ Plus TUNEL reaction was performed by incubation with the TUNEL reaction cocktail for 30 min at 37°C. Subsequently, the tissues were incubated with appropriate primary and secondary antibodies as described above. For detection of α2-6-linked sialic acid in pancreata, tissue sections were blocked by incubation with 5% BSA in 1X SiaFind™ Binding Buffer 2 (Lectenz Bio, BA0102) for 1 hour at RT, and then incubated with 25 μg/mL of SiaFind™ α2,6-Specific Reagent SureLight® 488 (Lectenz Bio, SP2602F) and Pancreatic Alpha Amylase (Abcam, ab189341) in 1X binding buffer with 5% BSA, overnight at 4°C. Slides were then washed in TBS with 0.05% Tween 20 (TBS-T) and incubated with Donkey anti-Sheep Secondary Antibody, Alexa Fluor™ 647 (Thermo Fisher Scientific, A-21448) for 1 hour at RT. Images were captured using a Nikon 80i Eclipse microscope for all IF staining of pancreatic tissues.

#### IF staining of organoids

Organoids were grown in 8-well chamber slides (Thermo Fisher Scientific, 154941) and treated with 100 ng EGF or TGFα for 5 min. Organoids were fixed with 4% paraformaldehyde in TBS for 15 min at RT, permeabilized by incubation in 0.5% Triton X-100 in TBS for 20 min at RT, and then blocked by incubation with 2% BSA in TBS for 1 hour at RT. Organoids were subsequently incubated overnight at 4°C with antibodies against ST6GAL1 (R&D Systems, AF5924), EGFR (Abcam, ab52894), phospho-EGFR (pTyr-1068, Abcam, ab40815), p44/42 MAPK (ERK1/2) (CST, 4695), phospho-p44/42 MAPK (p-ERK1/2) (Thr202/Tyr204, CST, 9101), AKT (CST, 4691), and phospho-AKT (Ser473, Abcam, 9271) overnight at 4°C. Organoids were then incubated with secondary antibody for 1 hour at 37°C followed by incubation in Hoechst solution (Thermo Fisher) for 5 min. Organoids were mounted using VECTASHIELD Vibrance® Antifade Mounting Medium (Vector, H-1700) and images were captured using an Olympus X-6000 confocal microscope. For quantification of fluorescent intensity, at least 5 separate organoids for each group were selected randomly, and using an Olympus FV10-ASW (version 3.1), fluorescent intensity was measured for phospho-EGFR, EGFR, phospho-ERK, ERK, phospho-AKT, and AKT. Normalized fluorescent intensity was calculated by dividing target intensity by Hoechst intensity for each field of view. The normalized phosphorylated targets were then divided by the respective normalized total targets to obtain a ratio representing the activated proportion of each signaling molecule. At least two independent experiments were performed for each signaling target.

### Immunoblotting

For immunoblots testing activation of EGFR, HPNE cells were treated with 50 ng/ml EGF or TGFα at varying time points. For immunoblots testing upregulation of anti-apoptotic proteins, HPNE cells and organoids were treated with EGF or TGFα (HPNE: 50 ng/ml EGF or TGFα; Organoids: 100 ng/ml EGF or TGFα) for 24 hours. Some samples were pre-treated with an ERK1/2 inhibitor (SCH772984, 100 nM) 1 hour prior to experiment. Treated cells were immunoblotted for the Bcl-2 family anti-apoptotic proteins, Bcl-xL and Mcl-1.

HPNE cells were lysed in RIPA buffer (Thermo Fisher Scientific, 89901) containing Halt™ protease and phosphatase inhibitors (Thermo Fisher Scientific, 78440). For organoid lysis, organoids were scraped into PBS with 0.5 mM EDTA and centrifuged at 300g for 5 min. Supernatant was aspirated and organoids were then incubated at 37°C in 0.05% Trypsin (Thermo Fisher Scientific, 25300054) for 15 min with periodic manual disruption by pipetting. Organoids were then lysed in RIPA buffer as described above. Lysate protein concentration was quantified by BCA (Thermo Fisher Scientific, 23225). Following SDS-PAGE, proteins were transferred to PVDF membranes. Membranes were blocked in 5% nonfat dry milk and incubated overnight at 4°C with antibodies against EGFR (CST, 4267), phospho-EGFR (Tyr-1068, CST, 3777), p44/42 MAPK (ERK1/2) (CST, 4695), phospho-p44/42 MAPK (p-ERK1/2) (Thr202/Tyr204, CST, 9101), AKT (CST, 4691), phospho-AKT (Ser473, CST, 9271), Mcl-1 (CST, 5453), Bcl-xL (CST, 2764), cleaved Caspase-8 (Asp387, CST, 8592, mouse specific), cleaved Caspase-8 (Asp374, CST, 9496, human specific), cleaved Caspase-3 (Asp175, CST, 9661), ST6GAL1 (R&D Systems, AF5924), and β-actin (CST, 5125). Membranes were then incubated for 1 hour in secondary antibody at RT and developed using Clarity Western ECL substrate (Bio-Rad). Images were generated and recorded with Odyssey M (LICORbio).

### Analyses of kinase activity

#### Kinomics assay

Kinomics experiments were performed by the Kinome Core at the Department of Radiation and Oncology at the UAB Heersink School of Medicine. HPNE EV and OE cell lysates were generated using M-PER™ Mammalian Protein Extraction Reagent (Thermo Fisher Scientific, 78501) with Halt™ protease and phosphatase inhibitors (Thermo Fisher Scientific, 78440). Lysates were analyzed on the Protein Tyrosine kinase (PTK) array and the Serine/Threonine kinase (STK) array using 15 µg (PTK) or 2 µg (STK) protein per a standard kinomic protocol. Phosphorylation data was collected over multiple computer-controlled pumping cycles, and exposure times (10-200ms) for ∼144 substrates per array. Raw image analysis was conducted using Evolve2, and comparative analysis, and upstream kinase prediction was done in BioNavigator v6.2 using scoring from Kinexus (www.phosphonet.ca). Raw reads per cycle were recorded with Barcode, Array, Sample Name, Cycle, and Exposure Time. After kinetic reads, postwash captures were done at 10, 20, 50, 100, and 200 ms exposures. Whole Chip comparative analysis (BioNavigator Upkin PTK v8.0/STK v6.0) was conducted between groups, generating Mean Final Scores (MFS; non-directional combined specificity and sensitivity) and Mean Kinase Statistic.

#### RTK Array

A human Phospho-Receptor Tyrosine Kinase (RTK) Array Kit (R&D Systems, ARY001B) was used to assess RTK activation. Briefly, 1.5 mL of 150 µg of HPNE EV and OE cell lysates were added to the human phospho-RTK array membranes and incubated overnight at 4°C. The membranes were subsequently incubated with an anti-Phospho-Tyrosine-HRP detection antibody solution for 2 hours at RT and developed with Chemi Reagent Mix. Signals from each RTK were quantified by densitometry using ImageJ (NIH).

### Proliferation and cell survival assays

#### Proliferation

For the 5-day cell proliferation assays, 5000 cells were seeded per well in a 96-well plate. Cell number at each day was quantified using the CyQUANT™ Cell Proliferation Assay Kit (Thermo Fisher Scientific, C7026). To determine the percentage of cells in S-phase under normal and low serum conditions, the Click-iT® EdU Flow Cytometry Assay Kit (Thermo Fisher Scientific, C10420) was used. Briefly, cells were cultured in either full media (5.0% FBS), or serum-depleted media (0.5% FBS) for 24 or 72 hours and cells were then incubated with 10 μM of EdU for 1.5 hours. Cells in S-phase were detected with the LSRFortessa™ Cell Analyzer (BD Biosciences) using 488 nm excitation with a green emission filter (530/30 nm).

#### Cell stressor assays

To test cell survival under low seeding density conditions, cells were seeded at 500 cells per well in a 6-well plate and allowed to grow for 3 weeks in media with either 5.0% FBS or 0.5% FBS. Cells were stained with crystal violet and then solubilized in 10% acetic acid. Relative cell number was determined by measuring absorbance at 590 nm. To test anchorage-independent survival, a soft agar colony formation assay was used. 5000 cells were seeded in a top layer of 0.6% agar and placed on top of a 1.0% agar layer in a 6-well plate. Cells were cultured for 3 weeks and colonies were quantified on 4 randomly chosen fields.

### Cell death assays

For TNFα-induced apoptosis, HPNE cells were pre-treated with 10 µM of cycloheximide (Selleckchem, S7418) for 1 hour and then treated with 100 ng/mL of recombinant human TNFα (R&D Systems, 210-TA) for 24 hours. Organoids were treated with 100 ng/mL of recombinant mouse TNFα (R&D Systems, 410-MT) for 24 hours. For FasL-induced apoptosis, HPNE cells were treated for 24 hours with 10 ng/mL of recombinant human FasL (R&D Systems, 126-FL) along with 10 µg/mL of cross-linking anti-polyHistidine Monoclonal Antibody (R&D Systems, MAB050). Organoids were treated for 24 hours with 10 ng/mL recombinant mouse FasL (R&D Systems, 526-SA) and 10 µg/mL of cross-linking antibody anti-polyHistidine Monoclonal Antibody (R&D Systems, MAB050). Media from the cultures was collected and used to measure LDH release using the CyQUANT™ LDH cytotoxicity assay (Fisher, C20300). To measure caspase activity, HPNE and organoid cultures treated for 24 hours with TNFα or FasL as described above were lysed and immunoblotted for cleaved caspase 3 and cleaved caspase 8.

### GEM Models

C57BL/6 mice expressing ST6GAL1 under the Rosa-26 gene promoter (*LSL-ST6GAL1*) were generated by the UAB Transgenic Mouse Facility as described previously (38). Briefly, the human ST6GAL1 CDS was amplified from a plasmid containing the *ST6GAL1* gene (Origene) and inserted into pRosa26-DEST (Addgene #21189), which contains an LSL (lox-STOP-lox) cassette. These mice were crossed to *Pdx1-Cre* mice (Jackson Laboratory, B6.FVB-Tg(Pdx1-cre)6Tuv/J) to generate “SC” mice with expression of the ST6GAL1 transgene in the pancreas. For the WT controls, *LSL-ST6GAL1* littermates (lacking Pdx1-Cre) were used. We previously verified expression of the ST6GAL1 transgene by IHC and confirmed increased α2-6 sialylation of acinar cells by staining with the SNA lectin and flow cytometry (32).

### Statistics

Statistical analyses were conducted using GraphPad Prism (version 9.5.1). ST6GAL1 expression in human pancreatic specimens was analyzed using the nonparametric Kruskal-Wallis test followed by Dunn’s multiple-comparison test. A nonparametric test was selected after evaluating the data with the D’Agostino-Pearson normality test. For other experiments, data were evaluated using a 2-tailed Student’s t test, 1-way ANOVA or 2-way ANOVA, with appropriate follow-up tests, where p < 0.05 was considered significant.

## Study approval

All animal studies were approved by the UAB Institutional Animal Care and Use Committee. Both males and females were included.

## Acknowledgments

This study was supported by NIH grant U01 CA233581 (SLB) and Department of Defense grants HT94252410441 (SLB), HT94252410014 (SI), and HT94252310306 (VD). MPM was supported by NIH postdoctoral fellowship grant F32 CA264906. BH was supported by the postdoctoral Relay for Life Champion of Mission Award by the American Cancer Society PF-24-1247682-01-IBCD. The authors gratefully acknowledge assistance from the following UAB Shared Facilities: Flow Cytometry and Single Cell Core Facility (supported by P30 AI027767; P30 CA013148; S10OD032296), UAB Tissue Biorepository, Tissue Procurement Shared Resource (supported by P30 CA013148), and Kinome Core.

