## Supplementary material for "ST6GAL1 sialyltransferase promotes acinar cell survival and tissue regeneration during pancreatitis": Figure_S1

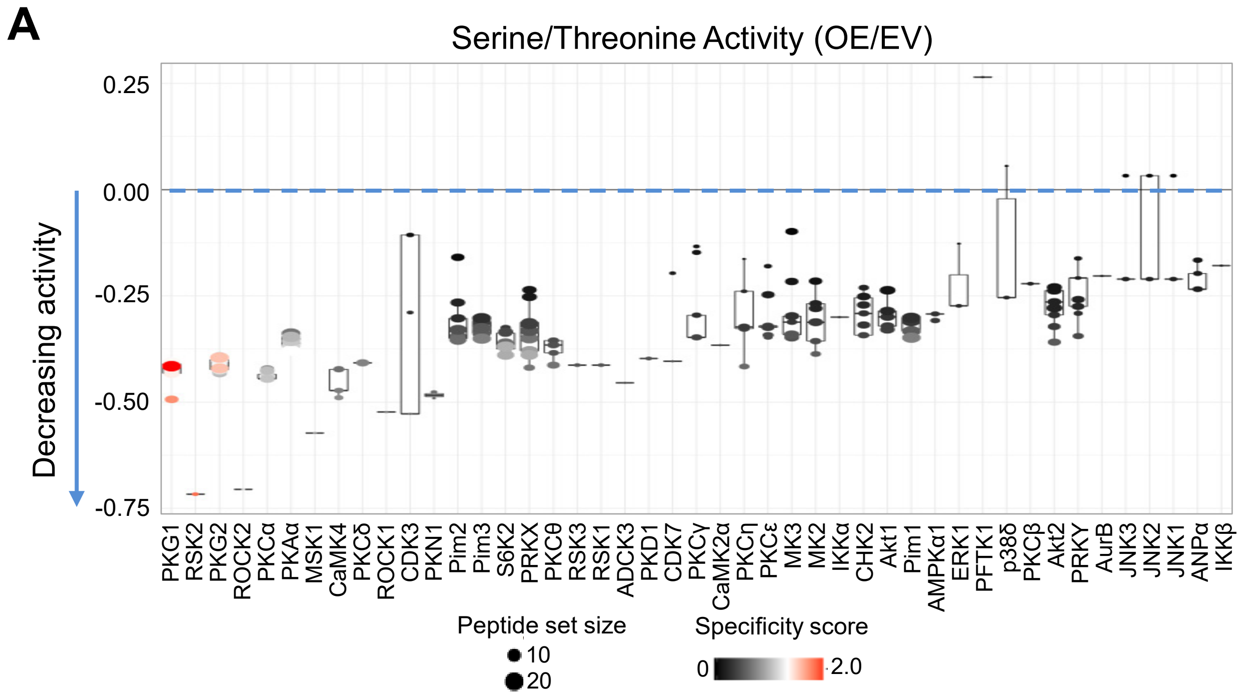


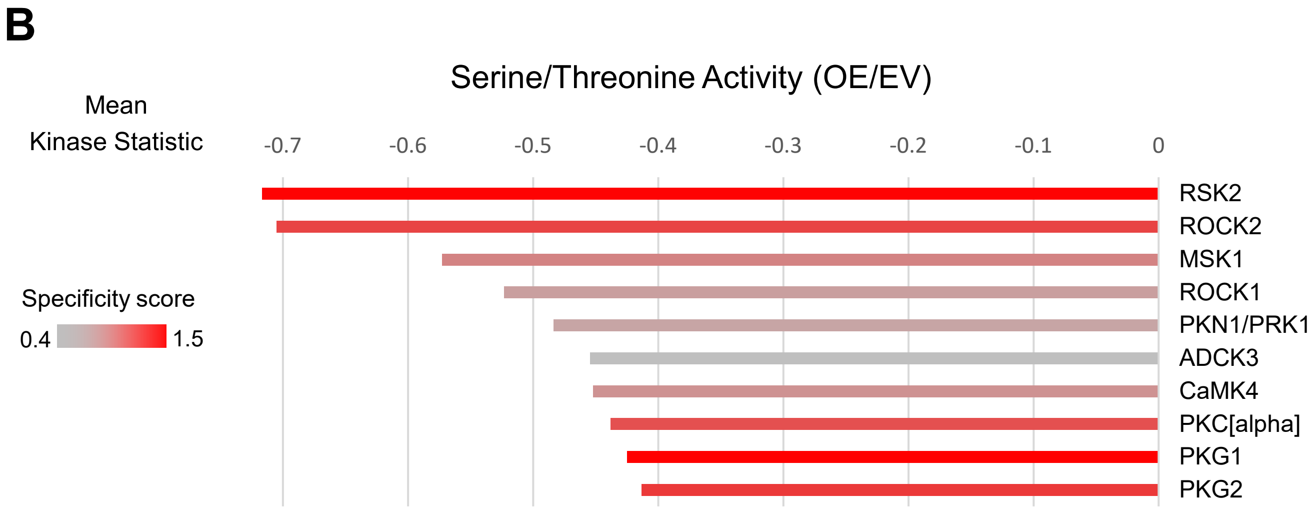


**Figure S1.** Serine/threonine kinase activity is reduced in cells with high ST6GAL1 expression. **(A)** Serine/threonine kinases identified using whole-chip (all peptide) comparison from the relative level of serine/threonine kinase activation in OE vs. EV cells. The Y-axis indicates the amount and direction of kinase change with a negative score indicating a decrease in kinase activity in OE cells. High specificity score indicates strong activity of the associated kinase on its intended target peptides. **(B)** Statistical values for the top 10 serine/threonine kinases decreased in OE vs. EV cells. The mean kinase statistic indicates the overall change in activity for a particular kinase.
